# Fast estimation of genetic relatedness between members of heterogeneous populations of closely related genomic variants

**DOI:** 10.1101/324418

**Authors:** Viachaslau Tsyvina, David S. Campo, Seth Sims, Alex Zelikovsky, Yury Khudyakov, Pavel Skums

## Abstract

Many biological analysis tasks require extraction of families of genetically similar sequences from large datasets produced by Next-generation Sequencing (NGS). Such tasks include detection of viral transmissions by analysis of all genetically close pairs of sequences from viral datasets sampled from infected individuals or studying of evolution of viruses or immune repertoires by analysis of network of intra-host viral variants or antibody clonotypes formed by genetically close sequences. The most obvious naϊeve algorithms to extract such sequence families are impractical in light of the massive size of modern NGS datasets. In this paper, we present fast and scalable k-mer-based framework to perform such sequence similarity queries efficiently, which specifically targets data produced by deep sequencing of heterogeneous populations such as viruses. The tool is freely available for download at https://github.com/vyacheslav-tsivina/signature-sj

## Introduction

Consider two sets *T*_1_ and *T*_2_ each containing *N* DNA or RNA sequences of length *L.* The *similarity join problem* consists in locating the set *P* of all pairs of sequences, with one sequence from *T*_1_ and the other from *T*_2_, within an edit distance or Hamming distance defined by the specified threshold *t*. In molecular epidemiology, this computational problem needs to be solved for detection of viral transmissions from sequences of intra-host viral variants sampled from infected individuals [1, 2]. Viral populations, for which the minimal inter-sample distance does not exceed the threshold, are considered to be potentially linked by transmission [1], while the number of pairs in *P* may suggest the time since a transmission event [3]. The related *genetic network construction* problem aims to build a graph with vertices corresponding to sequences from a given dataset *T* and edges corresponding to all pairs of sequences with an edit or Hamming distance less than the threshold *t*. This problem arises in studying and analysis of viral populations [4] or antibody repertoires [5]. Similar problems also emerged under different names in various areas of computer science [6, 7, 8, 9, 10].

The edit distance between a pair of sequences can be calculated in time *O*(*L*^2^) using dynamic programming [11]. If only distances below a desired threshold *t* which is small relative to *L* are desired. The distance calculation can be carried out with a small subset of diagonals neighboring the main diagonal of the dynamic programming matrix, leading to *O*(*tL*) time algorithm [12]. In this case a naϊve algorithm for the similarity join problem requiring pairwise comparison of all sequences has an asymptotic running time *O*(*tLN*^2^), which is still impractical for more than several thousand sequences.

Several *filtering-based approaches* have been put forward to improve the efficiency of the similarity join-type problems by reducing the number of pairs to be compared. Note that while fast heuristic and approximate methods exist such as Shingling[13], LSH[7], or BLAST[14], this paper focuses on the problem of exact distance calculation.

The common filtering approach is based on on the fundamental idea that related sequences should share long *k-mers* (substrings of length *k*) [15]. Several existing methods rely on signature schemes to quickly locate feasibly linked pairs [6] by assuming that pairs with an edit or Hamming distance which does not exceed a threshold *t* will share at least a certain number of *k*-mer-based signature keys. However, straightforward application of this technique to viral sequencing data is not sufficiently efficient, since mutations are not distributed uniformly along viral genomes, but tend to concentrate in short hypervariable regions [16]. As a result, many viral sequences share *k*-mers, thus significantly reducing the efficiency of filtering. The same effect has been observed for immunosequencing data [5], where all antibodies originating from the same V gene often share a *k*-mer from that gene.

In this paper, we describe a tool which uses k-mer-based signature filtering scheme optimized for viral data to solve the following problems:

- *Sample pair filtering:* given two NGS sequence samples *T*_1_ and *T*_2_, quickly determine whether the distances between all inter-sample pairs of sequences are greater than the threshold *t*.
- *Inter-sample sequence retrieval* (similarity join): given two NGS sequence samples *T*_1_ and *T*_2_, find all inter-sample pairs of sequences at edit distance or hamming distance below the threshold *t*.
- *Intra-sample sequence retrieval* (or genetic network construction): given an NGS sequence sample *T*_1_, find all pairs of sequences at edit distance or hamming distance below the threshold *t*.

The tool was validated using Hepatitis C Virus (HCV) data in the settings used for detection of viral transmissions and outbreaks [1, 2].

## 1 Methods

### 1.1 Notation

In the methods description, we assume that input sequence samples *T*_1_ and *T*_2_ both contain *N* sequences of length *L*, which cover the same genomic region. From here onwards we will use *k* as a fixed predefined parameter. Further we will use the following notation:

- *S* = *s*_1_*s*_2_ … *s*_*L*_ - sequence over the alphabet {*A, C, T, G*}.
- *S*[*i* : *j*] = *s*_*i*_*s*_*i*+1_… *s*_*j*_ - subsequence of *S* starting at position *i* and ending at position *j*.
- *k-mer* - any subsequence of length *k*
- *k-segment* - *k*-mer that starts at a position 1 + *ik, i* = 0,1, 2,….
- *K*(*S*) - the set of all *k*-mers of the sequence *S*.
- *R*(*S*) - the family of all *k*-segments of the sequence *S* (possibly with repetitions).
- *h*(*S*, *Q*) - Hamming distance between two sequences *S* and *Q*
- *l*(*S*, *Q*) - edit distance (Levenshtein distance) between two sequences *S* and *Q*
- 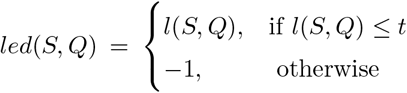 - limited edit distance, as mentioned above, could be calculated using dynamic programming [12]

### 1.2 Main Data Structure

Our signature-based filtering scheme is based on the following simple observation:

**Proposition 1.1** 
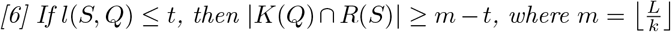

*Proof* If *S* and *Q* differ by an edit distance of *t*, then by the pigeon hole principal at most *t k*-segments differ between the sequences *S* and *Q*. So at least *m* – *t k*-segments must be the same.

Thus we need a fast way to calculate the number of common *k*-segments and *k*-mers for a given pair of sequences. To do it we introduce a hash function:

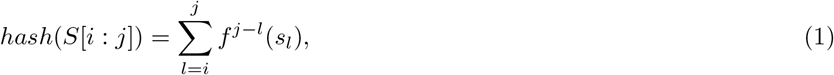

where *f* : {*A, C, G,T*} → {0,1, 2, 3} is an arbitrary bijection. For *k*-mers with *k* < 32, this hash function allows us to store them as 64-bit integers and can be quickly recursively calculated as follows:

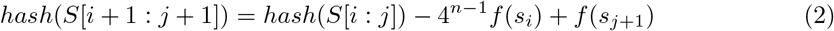

In addition, the hash can be inverted and so only the hash values of *k* – *mers* need to be stored.

In the proposed framework, each sample *T* is stored using a data structure further referred to as a *T*-dictionary and denoted by *dict*(*T*), which consists of the following fields:

- *dict*(*T*).*HM* - an inverted index of *T* [17], i.e. a hash table, where each key is a *k*-mer hash and its value is a set of all sequences from *T* that contain this *k*-mer.
- *dict*(*T*).*KM* - A set of all possible *k*-mer hashes in *T*
- *dict*(*T*).*KS* - hash table, where keys are sequences and values are lists of their *k*-segments (represented by their hash values) from 1 to *m*
- *dict*(*T*).*SC* - A list of *L* sets *SC*_1_,…, *SC*_*m*_, where *SC*_*i*_ is a set of all *k*-segments in a position 1 + *ik* (represented by their hash values).

### 1.3 Algorithm Description

We will first describe the approach for the sample pair filtering problem. Building a simple and fast filter for unrelated samples *T*_1_ and *T*_2_ is easy by applying Proposition 1.1 to whole samples as follows. Recall that *T*_1_ and *T*_2_ are considered to be genetically related, if the minimal edit distance between their sequences does not exceed the threshold *t*. Given two dictionaries *dict*(*T*_1_) and *dict*(*T*_2_), the necessary condition for their genetic relatedness is an existence of at least *m* – *t* positions {*i*_1_,*i*_2_,…, *i*_*m-t*_} such that *dict*(*T*_1_).*SC*_*ij*_ ∩ *dict*(*T*_2_).*KM* ≠ Ø for every *j* = 1, …, *m* – *t*. The sample pair filter pseudocode is presented at Algorithm 1.

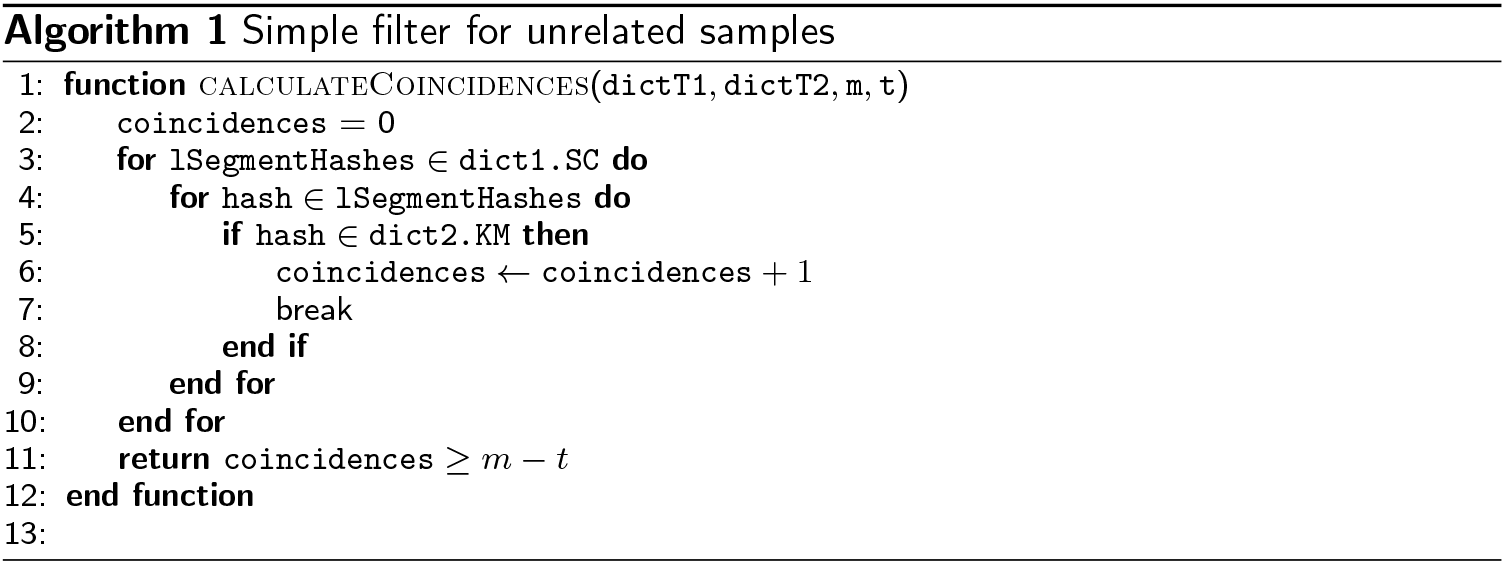

Assuming that membership verification for a hash set *dict*(*T*).*KM* can be performed in time *O*(1), the worst-case running time of the filter is *O*(*NL*). In real settings, samples with genetically related sequences produce significantly smaller maps *dict*(*T*).*SC*, thus leading to a lower average running time than in the worst case.

The algorithms for inter-sample sequence retrieval and intra-sample sequence retrieval problems are very similar, so we will describe the approach for the former problem. As before, let *T*_1_ and *T*_2_ be two samples. The algorithm first constructs the set of *candidate neighbors CN*_*S*_ ⊆ *T*_2_ for every sequence *S* ∈ *T*_1_. This procedure *(the filtering*), is followed by the *verification* procedure, which calculates actual neighbors of all sequences *S* ∈ *T*_1_ by calculating distances between *S* and all sequences *S*′ ∈ *CN*_S_. The pseudocode for inter-sample sequence retrieval algorithm is presented in Algorithm 2.

The basic filtering strategy utilizes Proposition 1.1, with the following features aiming at improvement of the running time. For each *S* ∈ T_1_, the set *CN*_S_ can be implemented as a hash table, with keys being sequences *S*′ ∈ *T*_2_ and values *CN*_S_(*S*′) being numbers of matches between *k*-segments in *S* and *k*-mers of *S*′. Let *L*_*S*_ be the number of *k*-segments of *S* that occur as *k*-mers in *T*_2_, and *I* = (*i*_1_, *i*_2_,… .*i*_*L*_*S*__) be the list of starting positions of these *k*-segments. To calculate the number of matches between *k*-segments in *S* and *k*-mers of *S*′ we may iterate over the list *I* and increment the current value of *CN*_*S*_(*S*′), when necessary. If after *j* iterations the inequality

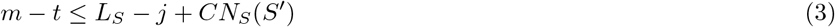

does not hold, then *S*′ cannot accumulate the required number of matches with the remaining iterations, and therefore the sequence *S*′ can be filtered out right away.

These considerations imply that the order in which starting positions of *k*-segments are examined is important in determining the algorithm’s running time.

The order of *k*-segment starting positions is determined heuristically as follows. For each position *i* let *k*_*S*_(*i*) = |*dict*(*T*_2_).*HM* (*S*[*i* : *i* + *k* − 1])| be the number of sequences from *T*_2_ that contain the *i*-th *k*-segment from *S*. If we sort positions by ascending order of the numbers *k*_*S*_(*i*) it usually leads to faster pruning of sequence pairs as this order minimizes the size of the candidate set that must be examined at each iteration.

Another simple adjustment could be implemented using the fact that the hamming distance is an upper bound for an edit distance, while the calculation of the former is significantly faster. Therefore if *h*(*S*, *Q*) ≤ *t*, then *Q* can be added to the list of neighbors of *S* without the edit distance calculation.

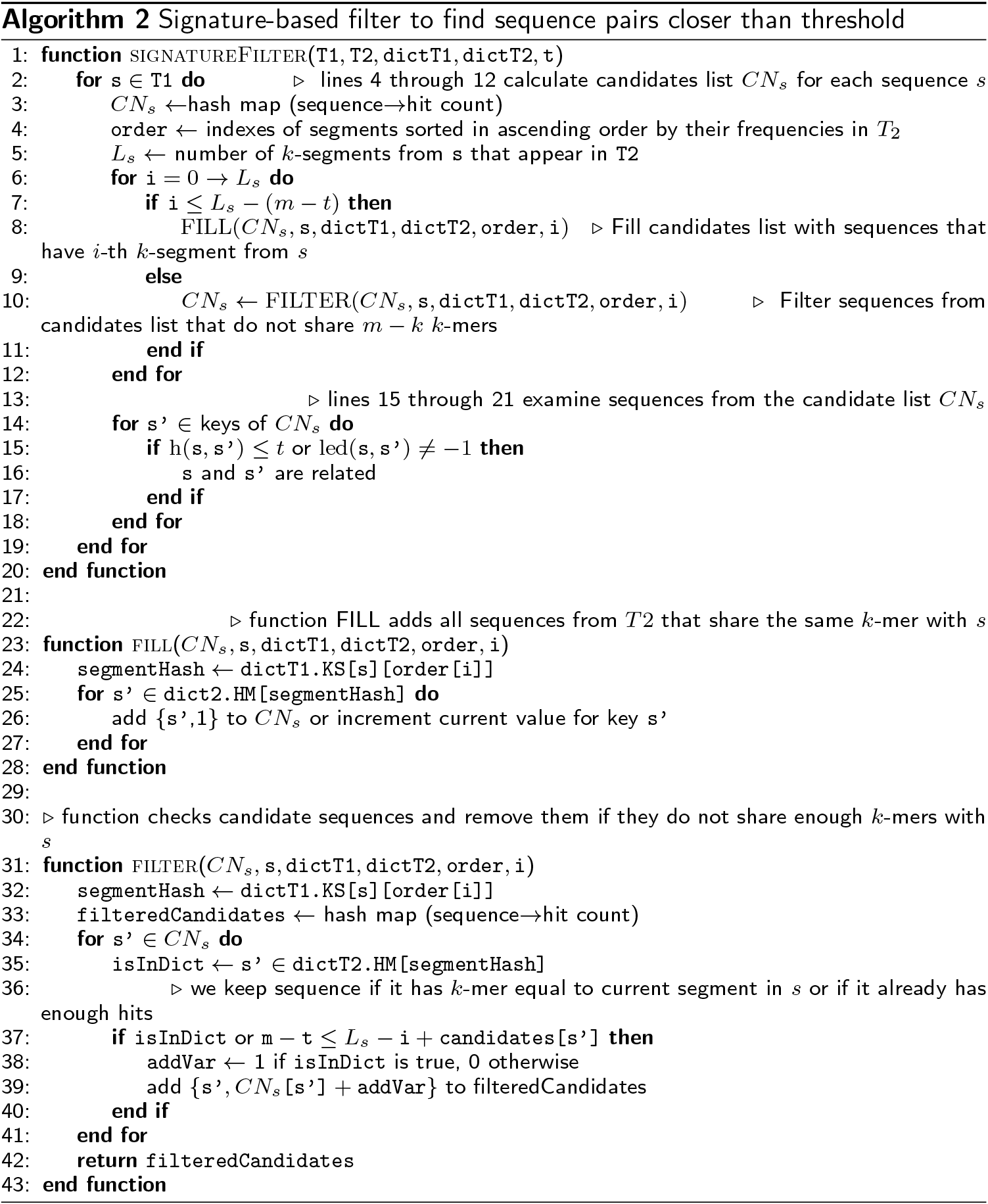

## 2 Hamming distance adjustment

The filtering strategy described above can be further improved, if the input sequences are aligned to a reference. In this case the samples can be compared using Hamming distance instead of an edit distance. For Hamming distance, Proposition 1.1 can be applied to *k*-segments of both comparable sequences thus simplifying the filling and filtering steps.

Furthermore, genomic heterogeneity is distributed highly irregularly along the genomes of species of interest. For example, Fig. 1 illustrates the distribution of nucleotide entropy for a particular intra-host population along the 264bp-long genomic HCV region at the junction of envelope glycoproteins E1 and E2, which is often used in epidemiological and immunological studies [18, 19, 1]. It should be noticed that *k*-segments from conserved regions are significantly less useful for the filtering as we want to maximize detectable differences between tested sequences. The non-uniformity in genomic heterogeneity can be taken into account by switching to the framework with *k*-segments of unequal size. By selecting *k*-segment boundaries that contain roughly equal amounts of average information entropy over the dataset, the filtering speed and quality could be significantly improved. Figure 6 provides an example, when entropy-based *k*-segments length allows more accurate filtering than uniform *k*-segments length.

**Figure 1.**
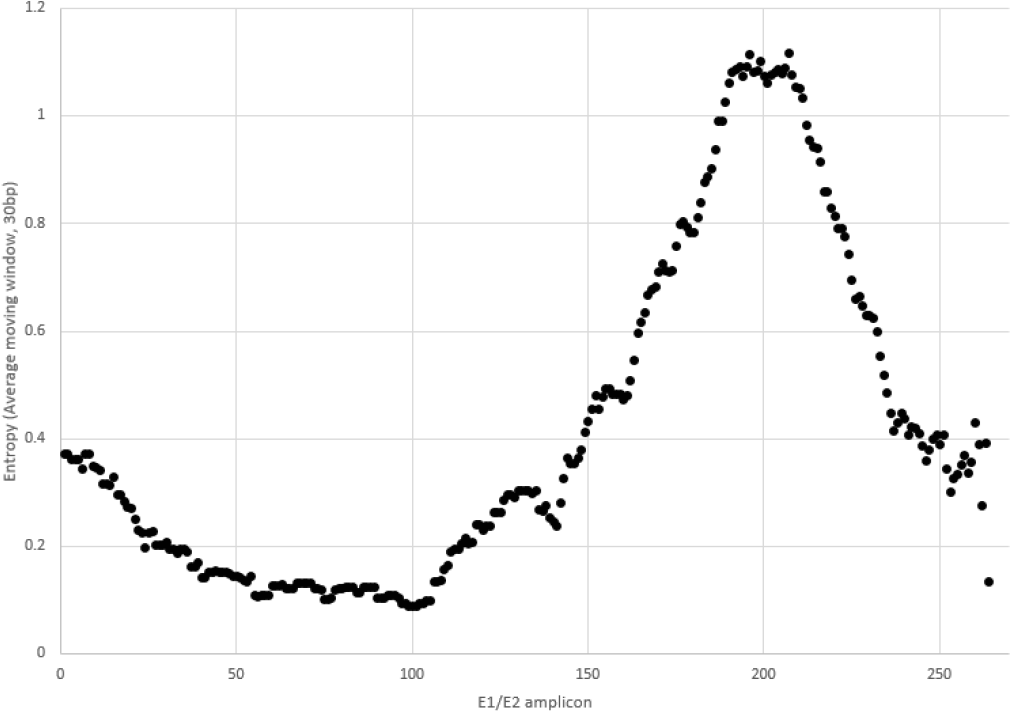
Distribution of nucleotide entropy along the E1/E2 region of HCV for a population of 469 unrelated genotype la sequences obtained from NCBI.

Formally, let 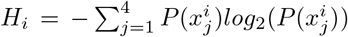 be the sample nucleotide entropy at position *i*, where 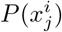 is a frequency of nucleotide 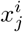 on *i*-th position of the alignment. The segments are selected in such a way that for every segment [*i*,*j*] we have 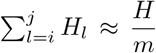, where 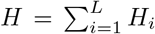 and *m* is the number of segments. Different numbers of segments were examined empirically and the best performance was obtained with *m* = *t* + 7.

## 3 Results

### 3.1 Validation Data

The developed tool was validated using NGS datasets of intra-host HCV populations sampled from infected individuals. Each dataset contains the E1/E2 junction of the HCV genome of length 264nt, which contains the Hyper Variable Region 1 (HVR1) region. Each sample was processed by error correction and haplotyping tools, and as a result we receive as an input datasets consisting of unique HCV haplotypes.

We used a set of 413 samples from [2] with 501.5 haplotypes per sample in average produced by NGS; 8 datatsets *d*_1_,…, *d*_8_ with 1000, 2000, …, 128 000 sequences produced by random sampling from NGS dataset with sequences sampled from chronically infected individuals and one additional NGS dataset *m*_1_ consisting of 10 467 sequences. The data are available in tool’s repository.

In all tests, the threshold *t* = 3.77% ≡ 10nt was used. This value is derived in [1] as empirically validated recommended threshold for separation between epidemiologically related and unrelated intra-host HCV HVR1 populations.

All tests were run on server with 128 Intel Xenon E7-4850 2.1GHz cores and 1.4Tb RAM. For Inter-sample sequence retrieval desktop PC was used with 4 Intel(R) Core(TM) i7-5500 2.4GHz cores and 8Gb RAM. All code is written on Java to provide a threaded, platform independent solution.

### 3.2 Sample pair filtering and Inter-Sample Sequence Retrieval validation

For Sample pair filtering and Inter-Sample Sequence Retrieval problems, we validated the tool using HCV datasets from [2]. The proposed approach has been compared with the Filter Composition pipeline proposed in [2]. Both methods were run on a desktop computer, as in the original paper [2]. The results are reported in Table 1. Here we show the result of comparison of all pairs of samples and all inter-sample pairs of sequences.

**Table 1.** Results of Filter Composition pipeline and *k*-mer based signature scheme filtering for Sample pair filtering and Inter-Sample Sequence Retrieval problems

| Method | Filter Composition | Signature Scheme |
| --- | --- | --- |
| Percent of filtered sample pairs | 85.1% | 92% |
| Percent of filtered sequence pairs | 91.5% | 99.996% |
| Total Time | ~ 5 min | ~ 15 sec |

The proposed sample pair filtration algorithm removed 92% of all possible samples pair comparisons, and sequence pair filtering algorithm managed to filter out 99.996% of all possible sequence pairs. The latter means that only 888,914 out of 22,037,502,011 sequence pairs passed from filtering to verification stage of the algorithm. As a result, the proposed approach significantly outperforms the Filter Composition Pipeline in filtering quality and in running time.

We studied how the filtering quality is affected by different optimization subroutines (Table 2). Disabling sample pair filtering increases the running time for comparison of all samples by 42%, while the impact of sorting of *k*-segment starting positions is even higher, with disabling of this step slowing down the comparison by 254%.

**Table 2.** Algorithm run time without optimization subroutines

| Feature | Time |
| --- | --- |
| No sample pair filter | ~ 21.3s |
| No sorting of $k$-segment starting positions | ~ 38.1s |

Preprocessing and dictionary building can take up a significant portion of the total running time of a signature-based filtering algorithm, when samples are distant and few distance calculations are required. For the given collection of 413 samples, preprocessing of all samples takes ~ 4840ms, which constitutes approximately 1/3 of the total running time of the algorithm. Note that in the case when significant number of closely related sequence pairs is present, the situation is different (see the next section).

The algorithm performance depends on the size of the *k*-mers and *k*-segments. Small *k* leads to larger number of matches between *k*-segments and *k*-mers of distant sequences, which can cause extra sequences to be added to the candidate lists thus leading to decrease in filtering quality. Larger *k* leads to fewer false matches but unfortunately also a larger *k*-mer dictionaries. We examined different *k*-mer sizes to determine the optimal size for our datasets and found that *k* = 11 gives the best performance.

### 3.3 Intra-sample Sequence Retrieval Validation

For Intra-sample Sequence Retrieval Problem, we validated the proposed approach using datasets d1,…,d8,m1. First, for it was compared with a single-thread, brute force method with the worst-case complexity *O*(*N*^*2*^*Lt*), which performs pairwise comparison of all sequences and calculates limited edit distance using dynamic programming as described in [12]. The results are presented in Table 3.

**Table 3.**
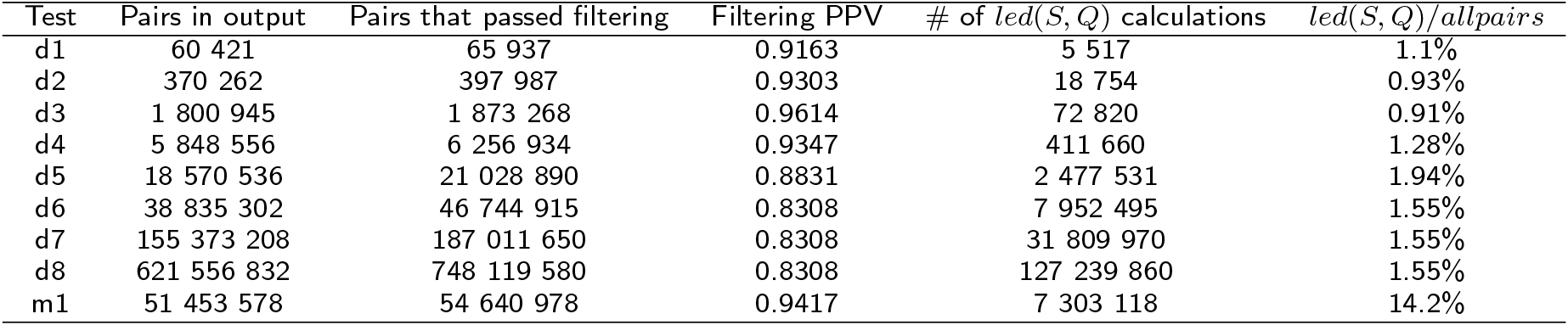
Intra-sample Sequence Retrieval Running Time

The running time of the proposed tool was also compared with the running time of a recently published method from [5], which was originally designed for the analogous problem for immunosequencing data. Fig. 2 illustrates that signature-based filtering approach demonstrates the significant advantage.

**Figure 2.**
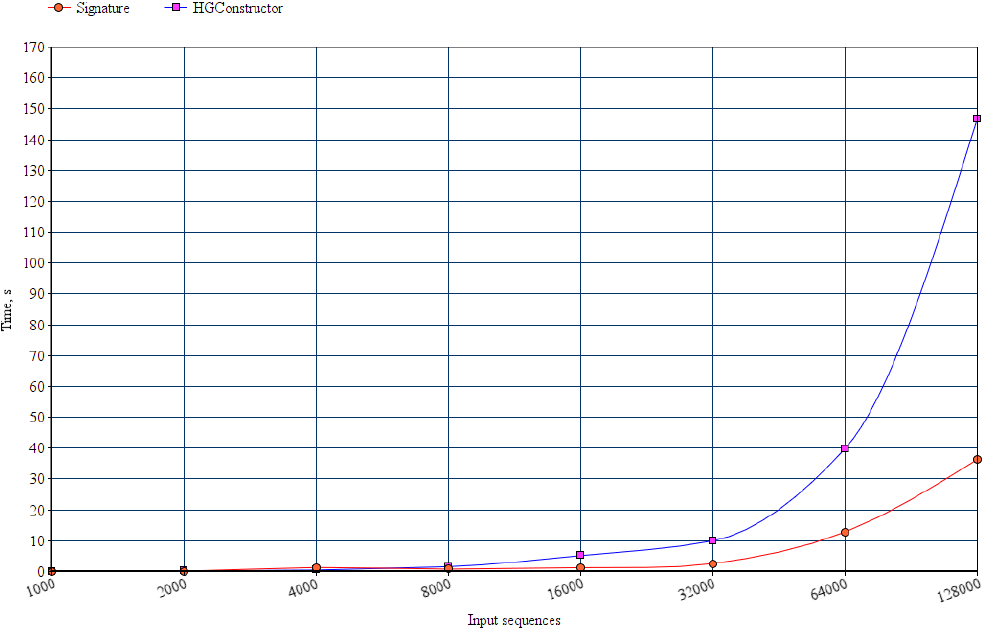
Running times of method from [5] (blue) and the proposed method (red) on datasets dl-d8

Fig. 3 demonstrates that for aligned sequences in most cases the adjustment utilizing entropy-based variable-size *k*-segments allows to achieve a significant speedup with respect to a constant-size *k*-segment.

**Figure 3.**
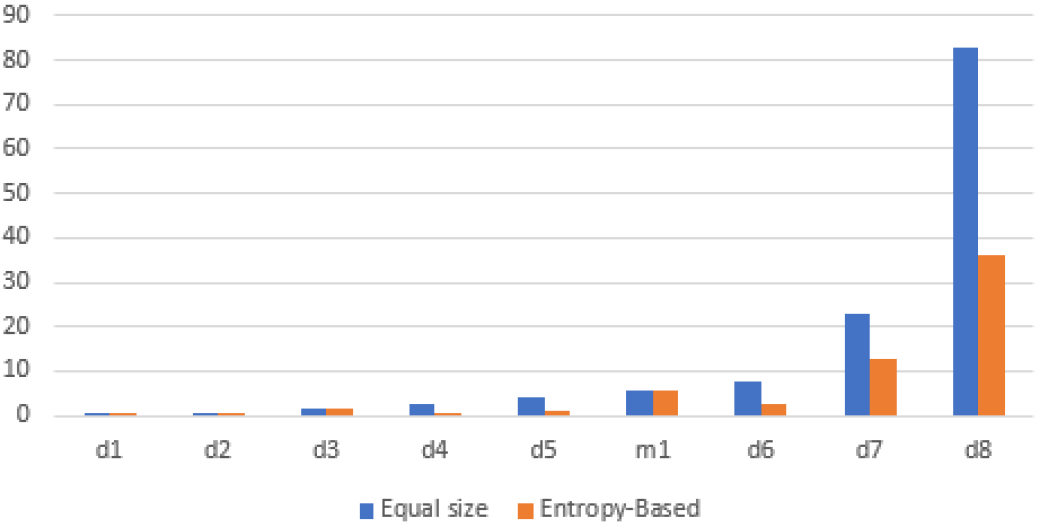
Comparison of running times of equal segment size and entropy-based approaches for single sample problem

The speedup described above is achieved by the combination of the several features. The first feature is the quality of filtering, which is analyzed in Table 4 and Table 5. On average, only ~ 10% of sequence pairs that pass filtering step (”false positives”) are not genetically related. As expected, most of the false positive pairs were very close to the threshold (Figure 4). With the threshold set at *t* = 10, pairs with an edit distance of *l*(*S*, *Q*) = 11,12,13 represent up to 75% of all false positives. Pairs that are so close to the threshold are difficult to filter out.

**Table 4.**
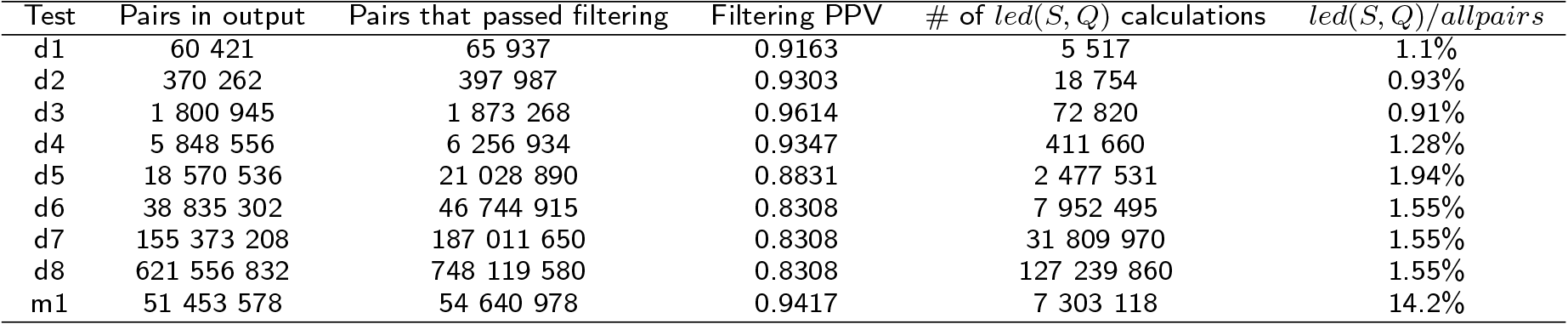
Filtering quality (unaligned sequences)

**Table 5.** Filtering quality (aligned sequences))

| Test | Pairs in output | Pairs that passed filtering | Filtering PPV |
| --- | --- | --- | --- |
| d1 | 60 420 | 64 573 | 0.9357 |
| d2 | 379 233 | 385 646 | 0.9834 |
| d3 | 1 800 448 | 1 862 914 | 0.9665 |
| d4 | 5 845 274 | 6 204 049 | 0.9422 |
| d5 | 18 551 359 | 20 706 813 | 0.8959 |
| d6 | 38 792 420 | 44 939 957 | 0.8632 |
| d7 | 155 201 680 | 179 791 828 | 0.8632 |
| d8 | 620 870 720 | 719 231 312 | 0.8632 |
| m1 | 47 101 270 | 48 888 011 | 0.9635 |

**Figure 4.**
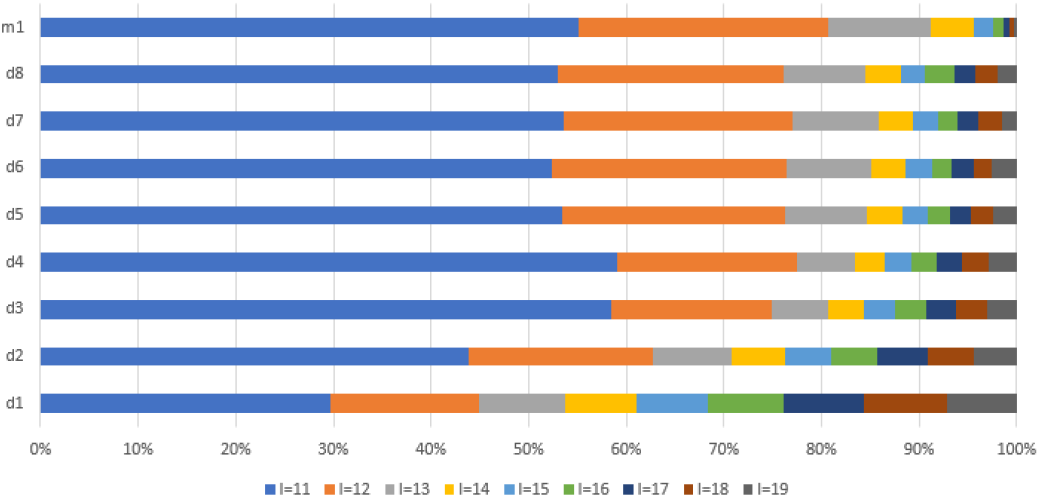
False positive sequence pairs(*l*(S,Q) > *t*) at different edit distances *l*

Another important feature is the fact that as the input increases in size the runtime of the algorithm is dominated by the edit distance calculations (Figure 5). However, the filtering and the Hamming distance shortcut reduces the number of edit distance calculations that must be performed. As a result, the actual edit distance is only calculated on small portion of the total pairs from the dataset (Table 4).

**Figure 5.**
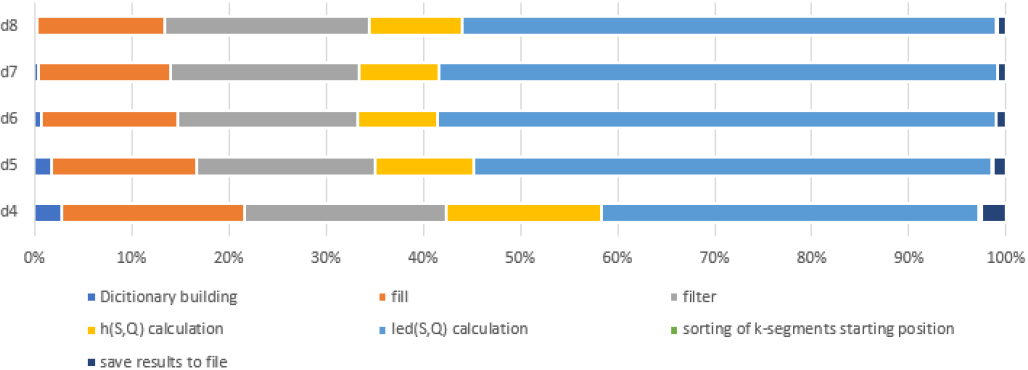
Contribution of algorithm subroutines to its total running time, unaligned sequences

We attempted to improve the filtering performance using other methods such as *k*-mer similarity [20], true matches [6], Hamming radius filter [2]. However, the overhead of these methods was greater than any runtime savings.

## 4 Conclusion

In this paper we presented an efficient signature-based tool to solve problems of edit or Hamming distance sequence retrieval for NGS data obtained from heterogeneous viral populations. It outperforms other approaches to this problem by including several data-specific steps and filters. The proposed approach was designed having problems of computational molecular epidemiology in mind. Until recent years, genomic analyses of viral transmissions and epidemic spread used a single viral sequence per infected individual. The advent of sequencing technologies now allows to analyze thousands of viral haplotypes per patient. Furthermore, just in the United States, from 2.7 million to 3.9 million people are infected with HCV [21], while ~ 1.1 million people are infected with HIV [22]. These numbers put an immense computational burden on real-time advanced molecular surveillance systems, such as Global Hepatitis Outbreak Surveillance Technology (GHOST)[23], which is currently being deployed by Centers for Disease Control and Prevention. When deployed, such system should have computational capacity to identify, whether a query set of viral samples is genetically related with any sample from a database consisting of hundreds of thousands of samples each consisting of thousands of sequences. The proposed approach aim to allow to process such queries efficiently. It builds on the general idea proposed in [6], which is heavily optimized by utilization of efficient data structures, such as inverted indexes and hash maps, and introduction of running time-improving procedures, such as efficient hash values calculation and determination of optimal order of *k*-mers processing. The proposed optimization steps allows for more than 2.5-fold running time decrease in comparison with the non-optimized filtering (Table 2). Furthermore, the proposed method takes into account uneven distribution of heterogeneous position along viral genomes by using variable entropy-based *k*-mers. It allows to improve both filtering quality (Fig. 6) and speed (Fig. 3). In general, for viral samples comparison the proposed filtering approach allows to eliminate the overwhelming majority of sequence comparisons and achieve a substantial running time decrease (Tables 1-5). The computational tool for efficient detection of genetic relatedness between genomic samples is freely available for download at https://github.com/vyacheslav-tsivina/signature-sj. The proposed tool should be especially useful for analysis of relatedness of genomes of viruses with unevenly distributed variable genomic regions, such as HIV and HCV. For the future we envision, that besides applications in molecular epidemiology the tool can also be adapted to immunosequencing and metagenomics data.

**Figure 6.**
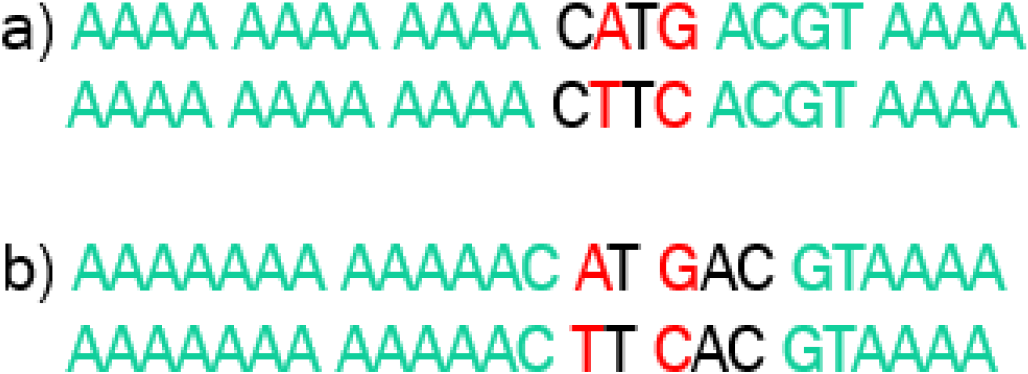
Example of two exact pairs of strings, but with equal (*k* = 4) (a) and entropy-based (b) segments size and *t* = 1. In case (a) the pair passes the filter, in case (b) it doesn’t pass the filter.

## Author’s contributions

VT designed, implemented and tested the algorithms and wrote the paper; DC and SS designed algorithms, prepared datasets for algorithms’ testing and wrote the paper; AZ designed algorithms; YK prepared datasets for algorithms’ testing, designed the study and supervised the research; PS designed the study, designed algorithms, prepared datasets for algorithms’ testing, wrote the paper and supervised the research.

## Acknowledgements

PS and AZ were partially supported by NIH grant 1R01EB025022-01

## References

1. Campo, D.S., Xia, G.-L., Dimitrova, Z., Lin, Y., Forbi, J.C., Ganova-Raeva, L., Punkova, L., Ramachandran, S., Thai, H., Skums, P., et al.: Accurate genetic detection of hepatitis c virus transmissions in outbreak settings. Journal of Infectious Diseases 213(6), 957–965 (2016)

2. Rytsareva, I., Campo, D.S., Zheng, Y., Sims, S., Thankachan, S.V., Tetik, C., Chirag, J., Chockalingam, S.P., Sue, A., Aluru, S., et al.: Efficient detection of viral transmissions with next-generation sequencing data. BMC genomics 18(4), 372 (2017)

3. Glebova, O., Knyazev, S., Melnick, A., Artyomenko, A., Khudyakov, Y., Zelikovsky, A., Skums, P.: Computational inference of transmission characteristics between viral populations. BMC Bioinformatics (accepted)

4. Skums, P., Zelikovsky, A., Singh, R., Gussler, W., Dimitrova, Z., Knyazev, S., Mandric, I., Ramachandran, S., Campo, D., Jha, D., et al.: Quentin: reconstruction of disease transmissions from viral quasispecies genomic data. Bioinformatics

5. Shlemov, A., Bankevich, S., Bzikadze, A., Turchaninova, M.A., Safonova, Y., Pevzner, P.A.: Reconstructing antibody repertoires from error-prone immunosequencing datasets. In: Research in Computational Molecular Biology, p. 396 (2017). Springer

6. Qin, J., Wang, W., Lu, Y., Xiao, C., Lin, X.: Efficient exact edit similarity query processing with the asymmetric signature scheme. In: Proceedings of the 2011 ACM SIGMOD International Conference on Management of Data, pp. 1033–1044 (2011). ACM

7. Gionis, A., Indyk, P., Motwani, R., et al.: Similarity search in high dimensions via hashing. In: VLDB, vol. 99, pp. 518–529 (1999)

8. Li, C., Wang, B., Yang, X.: Vgram: Improving performance of approximate queries on string collections using variable-length grams. In: Proceedings of the 33rd International Conference on Very Large Data Bases, pp. 303–314 (2007). VLDB Endowment

9. Medvedev, P., Scott, E., Kakaradov, B., Pevzner, P.: Error correction of high-throughput sequencing datasets with non-uniform coverage. Bioinformatics 27(13), 137–141 (2011)

10. Nikolenko, S.I., Korobeynikov, A.I., Alekseyev, M.A.: Bayeshammer: Bayesian clustering for error correction in single-cell sequencing. BMC genomics 14(1), 7 (2013)

11. Wagner, R.A., Fischer, M.J.: The string-to-string correction problem. Journal of the ACM (JACM) 21(1), 168–173 (1974)

12. Gusfield, D.: Algorithms on Strings, Trees and Sequences: Computer Science and Computational Biology, pp. 217–220. Cambridge university press, New York, NY, USA (1997)

13. Broder, A.Z., Glassman, S.C., Manasse, M.S., Zweig, G.: Syntactic clustering of the web. Computer Networks and ISDN Systems 29(8–13), 1157–1166 (1997)

14. Altschul, S.F., Gish, W., Miller, W., Myers, E.W., Lipman, D.J.: Basic local alignment search tool. Journal of molecular biology 215(3), 403–410 (1990)

15. Ma, B., Tromp, J., Li, M.: Patternhunter: faster and more sensitive homology search. Bioinformatics 18(3), 440–445 (2002)

16. Cuypers, L., Li, G., Libin, P., Piampongsant, S., Vandamme, A.-M., Theys, K.: Genetic diversity and selective pressure in hepatitis c virus genotypes 1-6: significance for direct-acting antiviral treatment and drug resistance. Viruses 7(9), 5018–5039 (2015)

17. Zobel, J., Moffat, A., Ramamohanarao, K.: Inverted files versus signature files for text indexing. ACM Transactions on Database Systems (TODS) 23(4), 453–490 (1998)

18. Pawlotsky, J.-M., Pellerin, M., Bouvier, M., Roudot-Thoraval, F., Soussy, C.-J., Dhumeaux, D.: Genetic complexity of the hypervariable region 1 (hvr1) of hepatitis c virus (hcv): Influence on the. Journal of medical virology 54, 256–264 (1998)

19. Bankwitz, D., Steinmann, E., Bitzegeio, J., Ciesek, S., Friesland, M., Herrmann, E., Zeisel, M.B., Baumert, T.F., Keck, Z.-y., Foung, S.K., et al.: Hepatitis c virus hypervariable region 1 modulates receptor interactions, conceals the cd81 binding site, and protects conserved neutralizing epitopes. Journal of virology 84(11), 5751–5763 (2010)

20. Peterlongo, P., Sacomoto, G.A.T., do Lago, A.P., Pisanti, N., Sagot, M.-F.: Lossless filter for multiple repeats with bounded edit distance. Algorithms for Molecular Biology 4(1), 3 (2009). doi:10.1186/1748-7188-4-3

21. Ward, J.W.: The hidden epidemic of hepatitis c virus infection in the united states: occult transmission and burden of disease. Topics in antiviral medicine 21(1), 15–19 (2013)

22. for Disease Control, C., Prevention, et al.: Diagnoses of hiv infection in the united states and dependent areas, 2015. HIV Surveillance Report 27, 1–114 (2016)

23. Longmire, A., Sims, S., Rytsareva, I., Campo Rendon, D., Dimitrova, Z., et al.: Ghost: Global health outbreak and surveillance technology. BMC Bioinformatics (accepted)

